# Development of SGC-GAK-1 as an orally active in vivo probe for cyclin G associated kinase through cytochrome P450 inhibition

**DOI:** 10.1101/629220

**Authors:** Christopher R. M. Asquith, James M. Bennett, Lianyong Su, Tuomo Laitinen, Jonathan M. Elkins, Julie E. Pickett, Carrow I. Wells, Zengbiao Li, Timothy M. Willson, William J. Zuercher

## Abstract

SGC-GAK-1 (**1**), a potent, selective, cell-active chemical probe for cyclin G associated kinase (GAK), was rapidly metabolized in mouse liver microsomes by P450 mediated oxidation. **1** displayed rapid clearance in mice, limiting its utility for in vivo studies. All chemical modifications of **1** that improved metabolic stability led to a loss in GAK activity. However, pretreatment of liver microsomes with the irreversible cytochrome P450 inhibitor 1-aminobenzotriazole (ABT) decreased intrinsic clearance of **1**. Coadministration of ABT also greatly improved plasma exposure of **1** in mice, supporting its use as a chemical probe to study the in vivo biology of GAK inhibition.

## Introduction

Cyclin G-associated kinase (GAK) is a 160 kDa member of the Numb-associated kinase (NAK) family of serine/threonine kinases.^1^ GAK was originally identified as a direct association partner of cyclin G and is ubiquitously expressed across tissues.^2^ Within the cell, GAK localizes to the Golgi complex, cytoplasm, and nucleus.^3^ GAK has been genetically associated with diverse range of biological processes. For example, genome-wide association studies have identified single nucleotide polymorphisms in *gak* that are associated with susceptibility to Parkinson’s disease.^4^ Knock down of GAK showed that it was required for maintenance of centrosome maturation and progression through mitosis.^5^ Conversely, GAK was found to be overexpressed in osteosarcoma cell lines and tissues where it contributed to proliferation and survival.^6^

The biological implications of GAK depletion or over-expression prompt the question of whether there is therapeutic utility in targeting its kinase domain with small molecules. Although GAK is a collateral target of several clinical kinase inhibitors, including erlotinib and gefitinib, until recently potent selective cell-active inhibitors of GAK had not been identified.^7^ Consequently, the biological consequences of selective GAK inhibition remained untested. As part of an ongoing effort to generate chemical probes for dark kinases, we recently described SGC-GAK-1 (**1**) as a high quality cell-active chemical probe for GAK (Figure 1).^8, 9^

GAK is a potential therapeutic target for prostate cancer. Expression of GAK is known to increase during prostate cancer progression to androgen independence and is positively correlated with Gleason score in resections from prostate cancer patients.^10^ We have demonstrated that **1** potently inhibits the viability of prostate cancer cells that contain constitively-active splice variants of the androgen receptor. These results prompted us to evaluate the effects of in vivo GAK inhibition in a prostate cancer xenograft model. We now describe our efforts to identify a chemical probe of GAK that is suitable for use in vivo.

**Figure 1.**
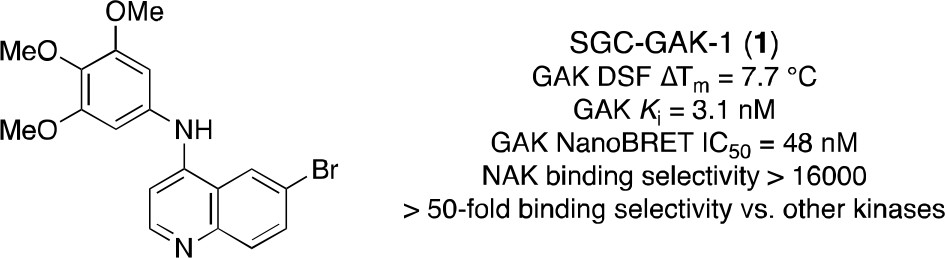
GAK cellular probe SGC-GAK-1 (**1**).

## Results

The metabolic stability of **1** was evaluated in mouse liver microsomes (MLMs) to assess the potential for extending the use of **1** from cellular to in vivo experiments. **1** was incubated with MLMs in the presence or absence of added NADPH, and compound was quantified at various time points over an hour. No degradation of **1** was observed in the absence of NADPH. However, **1** was rapidly consumed (T_1/2_ = 5.7 min; Cl_int_ = 990 mL/min/kg) in the presence of NADPH. This result suggested that **1** would be rapidly cleared in vivo and consequently be unlikely to attain sufficient pharmacokinetic exposure to be useful to study GAK in mouse models.

The necessity of NADPH for the metabolism of **1** was consistent with P450-mediated oxidation. To further understand the metabolic fate of **1**, we conducted a metabolite identification experiment. 10 μM of **1** was incubated for 5 min in MLMs at 37 °C in the presence of an excess of NADPH before quenching with methanol. After centrifugation, the supernatant was subjected to analysis by LCMS and the primary metabolite was identified as loss of CH_2_ from the 3,4,5-trimethoxyaniline, i.e. demethylated product **2** (Figure 2 and **Figure S1**). Minor metabolites were identified as an alternate loss of CH_2_ from the 3,4,5-trimethoxyaniline (e.g. **3**), the loss of 2 × CH_2_ (**4**), and a cyclic acetal (**5**). Formation of **5** likely occurs by the alcohol of a mono-demethylation product intercepting an intermediate oxocarbenium ion that would otherwise lead to the di-demethylated product. Finally, two minor species with quinoline ring oxidation (**6** and **7**) were observed at low abundance. The liver microsomal stability and the metabolite identification experiments identified cytochrome P450-mediated oxidation as the route of metabolism of **1**.

**Figure 2.**
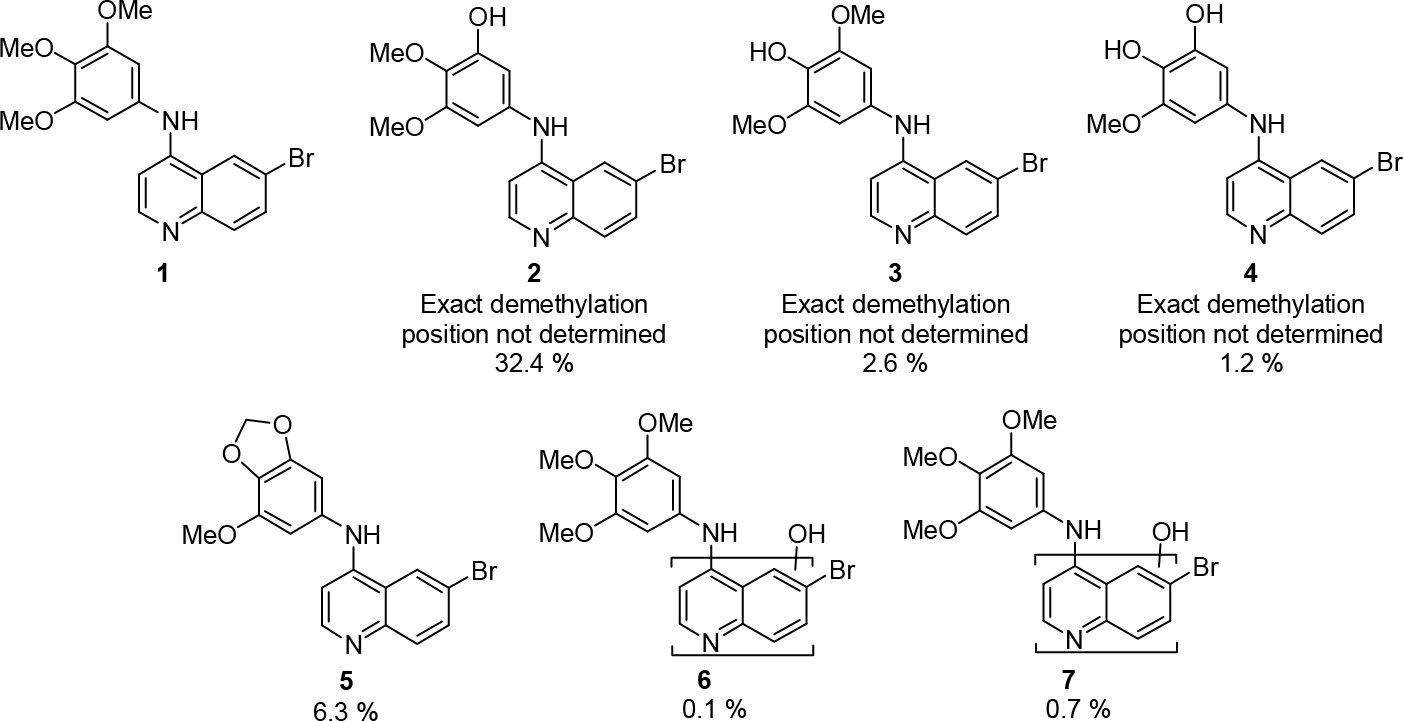
**1** and corresponding metabolites **2-7** identified along with percentage abundance with respect to parent compound. Compounds are ordered by HPLC retention time (see supporting information).

To further confirm that the metabolism of **1** was mediated by P450s we assessed the effect on compound stability of co-dosing with the broad spectrum P450 inhibitor 1-aminobenzotriazole (ABT).^11^ MLMs were pre-treated with 1 mM ABT for 30 min before exposure to 1 μM of **1**. The levels of **1** were quantified at various time points for 60 min. **1** showed a markedly improved stability in the presence of ABT (Table 1). The measured half-life was over 20 times longer, and the intrinsic clearance reduced to 44 mL/min/kg from 990 mL/min/kg. These results confirmed that **1** is primarily metabolized in liver microsomes by cytochrome P450-mediated oxidation.

**Table 1.** Microsomal stability assessment with and without ABT.

| Treatment | T<sub>1/2</sub> (min) | Cl<sub>int</sub> (mL/min/kg) |
| --- | --- | --- |
| **1** | 6 | 990 |
| **1** + ABT | 130 | 40 |
| Propanolol | 10 | 580 |
| Propanolol + ABT | 14 | 390 |

In an attempt to address the poor metabolic stability of **1**, we prepared a focused array of analogs. **8-11**, **13-31**, and **33-43** were synthesized through nucleophilic aromatic displacement of the corresponding 4-chloroquinolines (**Scheme 1**) to furnish the products in good to excellent yields (54-89 %).^8, 9, 12^ Additional analogs (**23**, **24**, and **27**) were synthesized by the same route in modest yields (12-38 %) due to the reduced nucleophilicity of their respective anilines. An alternative route employing a Buchwald-Hartwig cross-coupling with 4-chloro-6-trifluoromethylquinoline enabled access to the 6-trifluoromethyl analogs (**12**) and (**32**) in 21 % and 17 % yield, respectively (**Scheme 2**).^8, 9, 12^

**Scheme 1.**
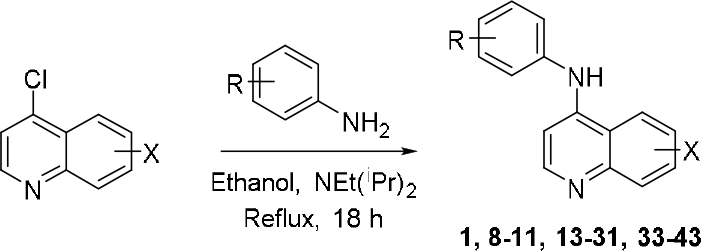
General displacement synthetic procedure

**Scheme 2.**
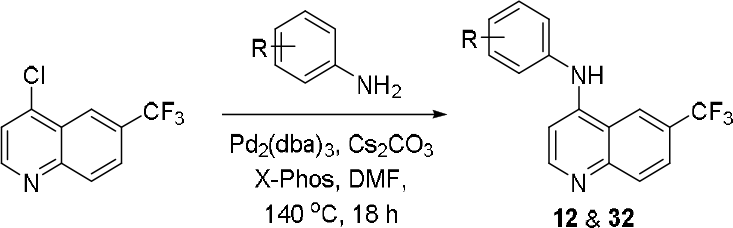
General Buchwald-Hartwig cross-coupling procedure

The metabolic stability of the 4-anilinoquinolines were evaluated in MLMs (Table 2). Half-lives were normalised to propranolol to control for experiments run on different days with different batches of liver microsomes. Changing the substitution of the core quinoline heterocycle to 6,7-dimethoxy as in **8** marginally increased stability relative to the 6-bromo substitution and maintained an aniline *O*-demethylation species as the primary metabolite. Modification of the aniline portion had a wide range effects on metabolic stability. The 2,3-methylenedioxy analog **9** showed similar stability to **8**, ith an aniline ring oxidation as the primary metabolite. Surprisingly, modification of the trimethoxy to the difluoro (**10** and **11**) yielded compounds that were equal to or less metabolically stable than **1**. Metabolite identification experiments with these fluorinated analogs continued to show aniline oxidation as the primary site of metabolism: direct ring oxidation was observed with **10** and hydroxyl replacement of a fluorine with **11**. Replacement of the trimethoxyaniline with a perfluorinated aniline as in **12** resulted in a significantly more stable compound, presumably due to complete deactivation of the aniline ring system. Finally, the stability of **13** suggested that 6-Br vs 6-CF_3_ quinoline substitution showed similar stability.

**Table 2.** Initial microsomal stability assessment.

| Compound | R | X | T<sub>1/2</sub> (min) | T<sub>1/2</sub> Ratio | Primary metabolism site | Primary metabolite mass |
| --- | --- | --- | --- | --- | --- | --- |
| **8** | 3,4,5-(OMe)<sub>3</sub> | 6,7-(OMe)<sub>2</sub> | 12.2 | 1.6 | Aniline | Parent – CH<sub>2</sub> |
| **9** | 2,3-O-CH<sub>2</sub>-O- | 6-Br | 8.9 | 1.9 | Aniline | Parent + O |
| **10** | 2,5-F | 6-Br | 1.7 | 0.17 | Aniline | Parent + O |
| **11** | 2,4-F | 6-Br | 3.2 | 0.47 | Aniline | Parent - F + OH |
| **12** | 2,3,4,5,6-F | 6-CF<sub>3</sub> | 45.6 | 7.0 | nt | nt |
| **13** | 3,4,5-(OMe)<sub>3</sub> | 6-CF<sub>3</sub> | 6.4 | 0.78 | Aniline | Parent + O |

nt: not tested

The analogs (**8-13**) were also assessed for GAK activity in a TR-FRET ligand binding displacement assay and a live cell NanoBRET target engagement assay (Table 3).^8, 13^ The 6,7-dimethoxyquinoline **8** displayed a high GAK cellular potency (IC_50_ = 25 nM) as did the methylenedioxy compound 9 (IC_50_ = 22 nM). The two difluoroaniline isomers **10** and **11** had low double digit nanomolar *K*_i_ values in the biochemical assay but only modest potency in the live cell NanoBRET assay. The metabolically stable pentafluoro aniline **12** had a significantly weaker GAK affinity (*K*_i_ = 880 nM) and was not progressed to evaluation in cells. Finally, **13** showed a 4-fold loss in cellular GAK potency relative to **1**. Considering both the metabolic stability and GAK affinity of these early compounds, we selected the methylenedioxy compound **9** for further optimization efforts.

**Table 3.** GAK affinity measurements and microsomal clearance of initial quinoline series.

| Cmpd | R | X | GAK K<sub>i</sub> (nM) | GAK IC<sub>50</sub> (nM) |
| --- | --- | --- | --- | --- |
| **8** | 3,4,5-(OMe)<sub>3</sub> | 6,7-(OMe)<sub>2</sub> | 0.54<sup>a</sup> | 25<sup>a</sup> |
| **9** | 2,3-O-CH<sub>2</sub>-O- | 6-Br | 1.9 | 22 |
| **10** | 2,5-F | 6-Br | 15 | 870 |
| **11** | 2,4-F | 6-Br | 35 | 2700 |
| **12** | 2,3,4,5,6-F | 6-CF<sub>3</sub> | 880 | nt |
| **13** | 3,4,5-(OMe)<sub>3</sub> | 6-CF<sub>3</sub> | 3.9<sup>a</sup> | 180<sup>a</sup> |

<sup>a</sup>Reference 11; nt = not tested.

Docking of **1** and **9** into a model of GAK derived from the cocrystal structure with gefitinib (Figure 3) was used to define key structural features for favourable binding using Schrödinger Maestro (Schrödinger Release 2018-4: LigPrep, Schrödinger, LLC, New York, NY, 2018).^14^ The compounds were minimised using LigPrep before docking into the ATP binding site prepared from PDB structure 5Y7Z.^15^ As expected, the most favourable docking pose placed the quinoline nitrogen in a position to accept a hydrogen bond from the backbone NH of the hinge region residue C126. The 5-membered ring of the methylenedioxy in **9** adopted a conformation where it could form a weak hydrophobic interaction with T123.^16^ Based on these results, we designed and prepared a series of isosteric replacements and positional isomers (**14-44**) that would maintain a favourable interaction with GAK and be less prone to oxidative metabolism. The analogs were assessed for GAK activity both biochemically and in cells and for MLM stability (Figure 4 and Table 4).

**Figure 3.**
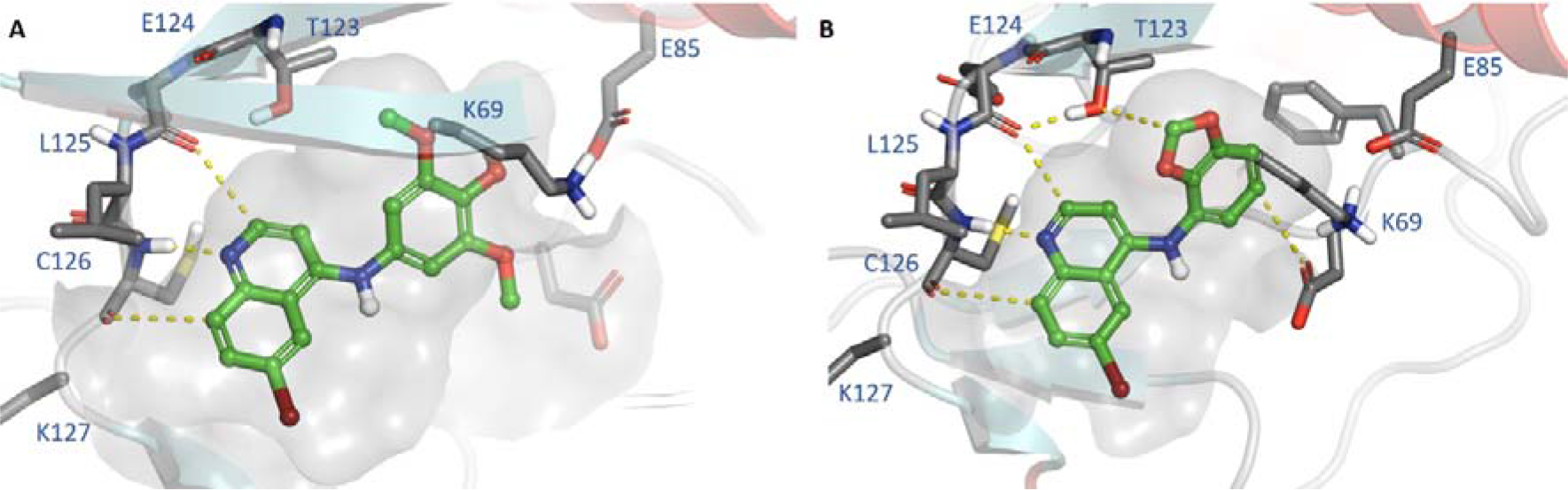
Docking of **1** (A) and **9** (B) into the ATP binding site of GAK (PDB: 5Y7Z).

**Figure 4.**
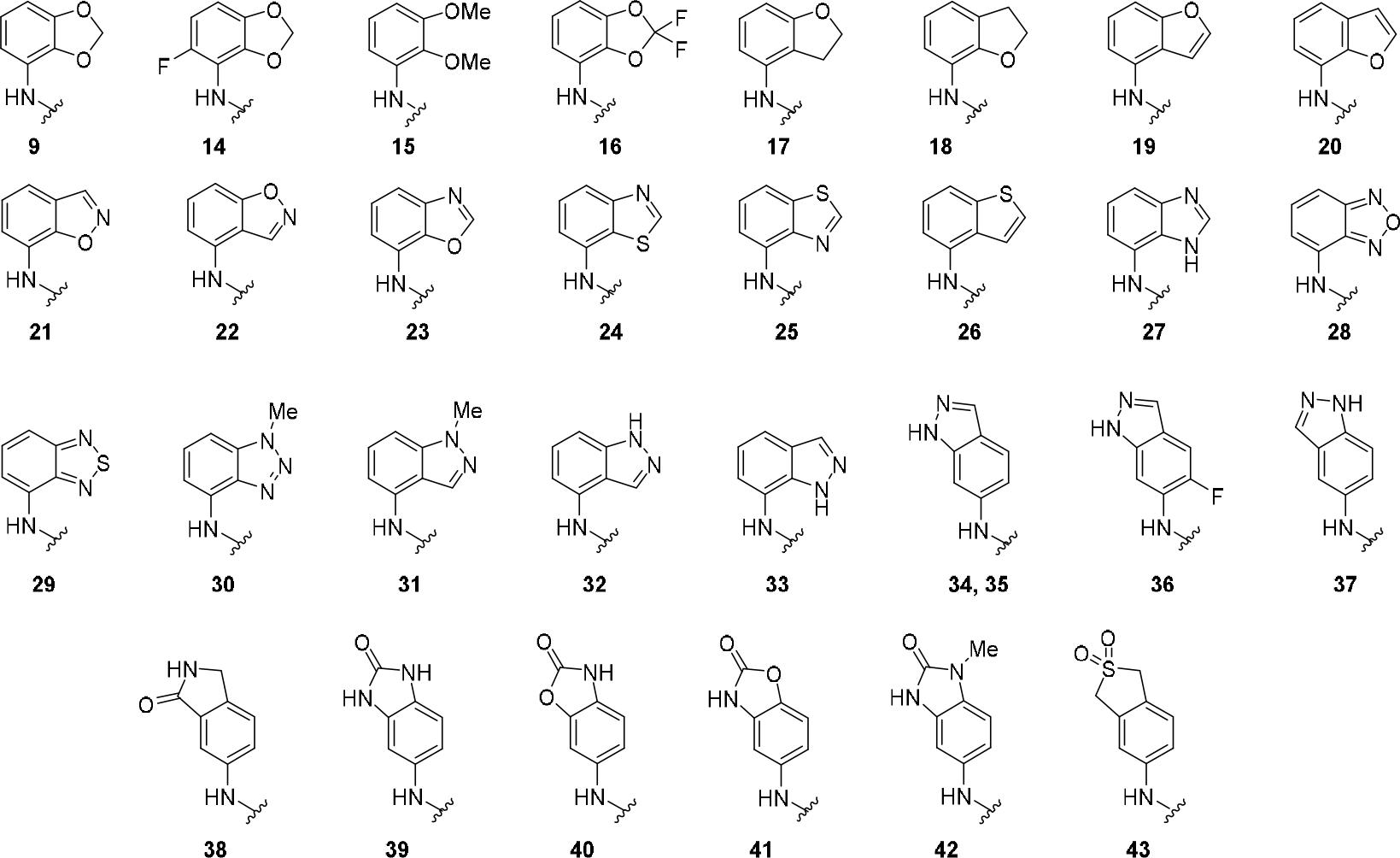
Aniline fragments incorporated into compounds **9** and **14 - 44** in Table 4.

Results of initial compound evaluation underscored the difficulty in concurrent optimization of GAK potency and metabolic stability (Table 4). Consistent with the docking model demonstrating a favorable fit for bicyclic aniline fragments, all of the analogs (**14-43**) demonstrated potent GAK activity in a biochemical assay (*K*_i_ < 100 nM) except the 2,3-dimethyoxy compound **15** (*K*_i_ = 740 nM) and the gem-difluoromethylenedioxy **16** (*K*_i_ = 460 nM). Unfortunately, potent biochemical GAK activity did not translate well to potent GAK activity in cells. Most of the heterocycles showed much lower GAK activity in the live cell NanoBRET assay compared to the methylenedioxy **9**. However, six compounds had submicromolar IC_50_ values in cells: benzofurans **19** and **20**, benzoxazole **23**, benzothiazole **24**, benzothiadiazole **29**, and indazole **32**. Interestingly switching to the 3,4-substitution from the 2,3-substitution often led to greatly increased metabolic stability with six compounds demonstrating T_1/2_ ratios ≥ 2-fold higher than **9**: indazoles **34** and **35**, benzimidazolone **39** and **42**, and benzoxazolidinones **40** and **41**. The benzimidazolone **39** was the most metabolically stable heterocycle with a T_1/2_ of > 200 minutes and T_1/2_ ratio > 25. Unfortunately, the matched pair of indazoles **34** and **35** were the only compounds to show any measurable activity in the live cell GAK NanoBRET assay with IC_50_ = 1.4 μM and 2.6 μM, respectively.

**Table 4.** GAK affinity measurements and microsomal clearance of isosteric replacements of **9**.

| Cmpd | X | GAK $K_i$ (nM) | GAK $IC_{50}$ (nM)<sup>a</sup> | $T_{1/2}$ (Min)<sup>a</sup> | $T_{1/2}$ Ratio |
| --- | --- | --- | --- | --- | --- |
| 9 | Br | 1.9 | 22 | 8.9 | 1.9 |
| 14 | Br | 1.4 | 120 | 1.8 | 0.27 |
| 15 | Br | 740 | >5000 | 1.0 | 0.12 |
| 16 | Br | 460 | >5000 | 2.0 | 0.24 |
| 17 | Br | 5.2 | >5000 | 1.7 | 0.16 |
| 18 | Br | 29 | 2500 | 0.82 | 0.10 |
| 19 | Br | 1.9 | 180 | 1.7 | 0.16 |
| 20 | Br | 4.3 | 570 | 0.9 | 0.08 |
| 21 | Br | 37 | 1400 | 5.2 | 0.64 |
| 22 | Br | 19 | >5000 | 1.4 | 0.17 |
| 23 | Br | 19 | 340 | 1.7 | 0.21 |
| 24 | Br | 11 | 220 | 5.3 | 0.68 |
| 25 | Br | 26 | >5000 | 2.6 | 0.32 |
| 26 | Br | 23 | 3300 | 4.1 | 0.50 |
| 27 | Br | 4.1 | >5000 | 8.1 | 1.0 |
| 28 | Br | 41 | 2500 | 1.2 | 0.15 |
| 29 | Br | 5.3 | 400 | 1.6 | 0.20 |
| 30 | Br | 2.8 | >5000 | 0.73 | 0.09 |
| 31 | Br | 3.6 | >5000 | 0.80 | 0.10 |
| 32 | Br | 14 | 760 | 2.5 | 0.32 |
| 33 | CF<sub>3</sub> | 720 | >5000 | 5.9 | 0.90 |
| 34 | CF<sub>3</sub> | 34 | 2600 | 46.4 | 9.8 |
| 35 | Br | 20 | 1400 | 45.6 | 9.8 |
| 36 | Br | nt | >5000 | 14.8 | 2.2 |
| 37 | Br | 47 | >5000 | 7.5 | 0.92 |
| 38 | Br | 62 | >5000 | 8.1 | 1.0 |
| 39 | Br | 360 | >5000 | > 200 | > 25 |
| 40 | Br | 680 | >5000 | 55.9 | 6.9 |
| 41 | Br | 110 | >5000 | 27.8 | 3.4 |
| 42 | Br | 1600 | >5000 | 33.6 | 4.2 |
| 43 | Br | 1700 | >5000 | 15.6 | 1.9 |

nt = not tested; <sup>a</sup> Mean average, n = 2

Although we were able to stabilise **1** by modification with an indazole head-group, we were ultimately unable to identify an analog that also maintained sub-micromolar potency in cells, a threshold we consider essential for chemical probes to be useful. Having been unable to identify a cell active GAK inhibitor with improved metabolic stability, we decided to test an alternative strategy to develop an in vivo chemical probe. Since in vitro inhibition of P450s by cotreatment with ABT in microsomes had greatly increased the metabolic stability of **1**, we decided to explore whether codosing would translate into in vivo stabilization. A pharmacokinetic experiment was performed in which C57BL/6 mice were dosed orally with 10 mg/kg of **1** with or without 2 h pre-treatment with 50 mg/kg ABT (Table 4). As predicted by the microsomal stability experiment, orally dosed **1** was rapidly cleared in mice with a T_1/2_ < 1 h and a low C_max_ (< 100 ng/mL). In contrast, pretreatment with ABT resulted in a dramatically extended T_1/2_ and more than 20-fold higher exposure. In the absence of ABT, the plasma concentration of **1** fell below the cellular IC_50_ after approximately 1 h. In contrast, pretreatment with ABT resulted in the sustained plasma concentration of **1** at two-fold above the cellular IC_50_ for 6 h with a 10 mg/kg dose (Figure 5). These results demonstrate the utility of co-dosed ABT in decreasing metabolism of **1** and provide a method for use of ABT-**1** as an in vivo chemical probe.

**Table 4.**
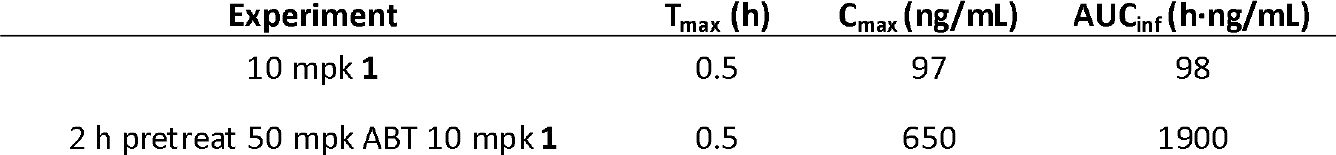
Pharmacokinetic profile of **1** with and without pre-administration of ABT.

**Figure 5.**
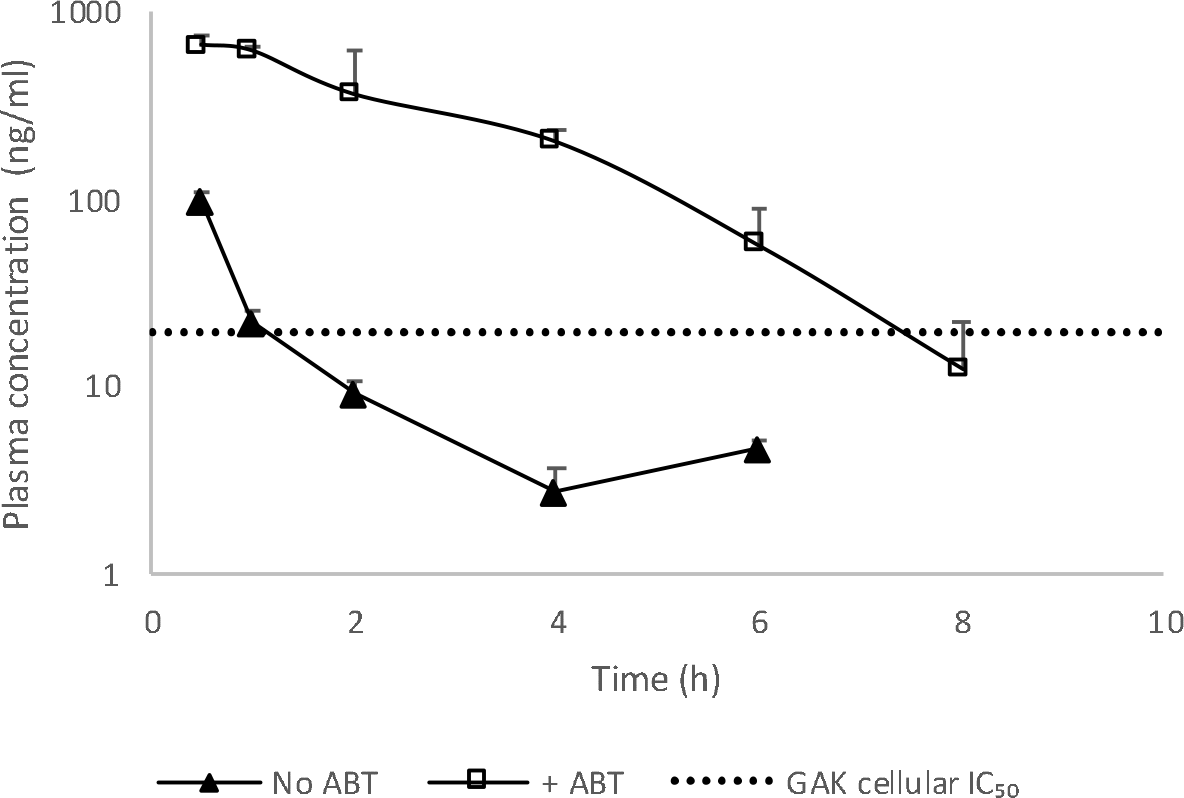
Plasma concentration profiles of **1** relative to cellular potency.

## Discussion

Chemical probes are potent and selective modulators of protein function useful as reagents to elucidate the biology of their targets.^17, 18^ A significant amount of learning can be derived from use of chemical probes in cells. Positive results in cellular experiments naturally lead to interest in using the chemical probe for target validation in animals. However, many chemical probes are not amenable to use in vivo because they are unable to attain sufficient exposure to support valid experiments. The pharmacokinetic profiles of probes can be addressed through careful chemical optimization of the series but at the cost of significant investment of time and resource.

A common pharmacokinetic liability of compounds is that they are rapidly metabolized by cytochrome P450 enzymes in the liver and rapidly cleared from the organism, a process known as first pass clearance. As a result, compounds are routinely evaluated for stability in MLM prior to progression to in vivo pharmacokinetic studies.^19^ It has been well established that molecules with low MLM stability are prone to rapid first pass clearance in vivo. The challenge of optimizing compounds to improve in vivo pharmacokinetic properties while maintaining their target potency and selectivity has vexed many medicinal chemistry programs, including our exploration of the dark kinase GAK.

1-Aminobenzotriazole (ABT) is a potent inhibitor of CYP3A4, CYP2D6, and other P450 enzymes, and it has been successfully employed to enhance the pharmacokinetics of metabolically labile compounds.^20^ For example, co-dosing of ABT with the antifungal APX001 increased exposure of the compound and enhanced its efficacy in a murine model of invasive candidiasis.^21^ Although not a viable strategy for improving exposure for medicines administered to human patients, co-dosing with a P450 inhibitor such as ABT or ritonavir can enable in vivo target validation experiments that would otherwise not be possible.^22^ The use of ABT to block oxidative metabolism is significantly less demanding of time and resource than chemical optimization and may be broadly applicable for development of in vivo chemical probes. For compounds metabolized by cytochrome P450 enzymes not inhibited by ABT (such as CYP2C9), other inhibitors such as atipamezole are available and have demonstrated utility in enhancement of pharmacokinetic profiles.^23^

The present report supports the utility of **1** and ABT in studying the in vivo biology of acute GAK inhibition in mice. Previous studies of ABT suggest the potential for broader applicability. First, longer term dosing of ABT is possible. For example, ABT displayed a well-behaved pharmacokinetic profile over 5 days dosing via osmotic pump.^24^ Minimal side effects were observed on chronic ABT dosing, the most prominent of which is gastric emptying.^25^ Second, there is potential for **1** and ABT to be utilized in species other than mice in in vivo studies. ABT is a competent cytochrome P450 inhibitor in rats, guinea pigs, and other species and has been used to enhance pharmacokinetic exposure in these contexts.^11, 26^

Although we were able to attain useful in vivo exposure through coadministration of ABT, our goal of chemical optimization of **1** remained unrealized. Replacement of the 3,4,5-trimethoxy substitution with 5-membered heterocycles annelated at the aniline 2,3-positions yielded several compounds that maintained high GAK affinity in a biochemical assay but without improvement in microsomal stability. On the other hand, compounds with 3,4-annelation of heterocycles in general proved to be significantly more metabolically stable but at the cost of weakened GAK affinity. The identification of heterocyclic replacements of the 3,4,5-trimethoxyphenyl group with increased microsomal stability is a valuable learning. The 3,4,5-trimethoxyphenyl group is a commonly encountered metabolically labile substructure in ATP-competitive inhibitors of protein kinases and in a variety of molecules with biological activity. It is a substructure in several approved drugs including the antibiotic trimethoprim, the antihypertensive reserpine, the anti-inflammatory colchicine, and the DHFR inhibitor trimetrexate for the treatment of leiomyosarcoma.^27^ Thus, future medicinal chemistry efforts involving optimization of the series with the 3,4,5-trimethoxyphenyl group may consider incorporation of select heterocyclic replacements identified here.

In conclusion, the cellular chemical probe SGC-GAK-1 (**1**) undergoes oxidative metabolism in mouse liver microsomes and is rapidly cleared in vivo. Pre-administration of ABT dramatically increases the pharmacokinetic exposure of **1** in mice and enables a sufficient exposure to support in vivo target validation experiments. The combination of **1** and ABT can be used as an in vivo GAK chemical probe in mice.

## Experimental

### Computational methods

Small molecules were prepared using ligprep module of Schrodinger (Schrödinger Release 2018-4: LigPrep, Schrödinger, LLC, New York, NY, 2018.) Structures were superimposed using flexible ligand alignment functionality (Schrödinger Release 2018-4: Maestro, Schrödinger, LLC, New York, NY, 2018) using compound **8** as template and employing Bemis-Murcko method for common substructure recognition.^14^

**NAK family TR-FRET assay** was performed as previously described.^12^

**GAK NanoBRET assay** was performed as previously described.^9^

**Metabolite identification**. compounds were incubated at a concentration of 10 μM with mouse liver microsomes, in the presence of an excess of NADPH. The duplicate incubations were performed in 0.5 mL 96-well plates in a shaking water bath maintained at 37 °C for 30 minutes and quenched by the addition of an equal volume of methanol. (see SI for full details and reports)

### Pharmacokinetic profiling

Pharmacokinetic profiling was performed by SAI Life Sciences. Healthy male C57BL/6 mice (8-12 weeks old) weighing between 20 to 35 g were procured from Global, India. Three mice were housed in each cage. Temperature and humidity were maintained at 22 ± 3 °C and 30-70 %, respectively and illumination was controlled to give a sequence of 12 hr light and 12 hr dark cycle. Temperature and humidity were recorded by auto controlled data logger system. All the animals were provided laboratory rodent diet (Envigo Research private Ltd, Hyderabad). Reverse osmosis water treated with ultraviolet light was provided ad libitum. 18 male mice were weighed and divided in to two groups as Group 1 and Group 2 with 9 mice in each group. Animals in Group 1 were administered orally with solution formulation of SGC-GAK-1 at 10 mg/kg dose. Animals in Group 2 were administered orally with solution formulation of ABT (2 hr before dosing of SGC-GAK-1) at 50 mg/kg dose. After 2 hr of ABT administration, animals were administered orally with solution formulation of SGC-GAK-1 at 10 mg/kg dose. Solution formulation of SGC-GAK-1 was prepared in 5% NMP, 5% solutol HS-15 in normal saline. The dosing volume administered was 10 mL/kg for oral route. Blood (~60 μL) samples were collected from set of three mice at each time point in labeled micro centrifuge tube containing K_2_EDTA solution as anticoagulant at pre-dose, 0.5, 1, 2, 4, 6, 8, 12 and 24 hr. Plasma samples were separated by centrifugation at 4000 rpm for 10 min and stored below −70 °C until bioanalysis. Concentrations of SGC-GAK-1 in mice plasma samples were determined by fit for purpose LC-MS/MS method. The plasma concentration-time data for SGC-GAK-1was provided for data analysis by bioanalytical group of Sai Life Sciences Ltd, Pune. The plasma, concentration-time data were then used for the pharmacokinetic analysis. Non-compartmental-analysis module in Phoenix WinNonlin^®^ (Version 6.3) was used to assess the pharmacokinetic parameters. Maximum concentration (C_max_) and time to reach the maximum concentration (T_max_) were the observed values. The areas under the concentration time curve (AUC_last_ and AUC_inf_) were calculated by linear trapezoidal rule.

### Chemistry

Compounds **1**, **8** and **13** were synthesized as previously reported.^8, 9, 12^ Compound spectra of **9-12**, **14-43** are included in the Supporting Information. All compounds were > 98% pure by ^1^H/^13^C NMR and LCMS.

### General procedure for the synthesis of 4-anilinoquinolines

6-Bromo-4-chloroquinoline (1.0 eq.), 3,4,5-trimethoxyaniline (1.1 eq.), and ^i^Pr_2_NEt (2.5 eq.) were suspended in ethanol (10 mL) and refluxed for 18 h. The crude mixture was purified by flash chromatography using EtOAc:hexane followed by 1-5 % methanol in EtOAc. After solvent removal under reduced pressure, the product was obtained as a free following solid or recrystallized from ethanol/water.

***N*-(Benzo[*d*][1,3]dioxol-4-yl)-6-bromoquinolin-4-amine (9)** was obtained as a yellow solid (157mg, 0.458 mmol, 74 %). MP > 270 °C decomp.; ^1^H NMR (400 MHz, DMSO-*d*_6_) δ 11.21 (s, 1H), 9.24 (d, *J* = 1.9 Hz, 1H), 8.57 (d, *J* = 6.9 Hz, 1H), 8.25 – 8.02 (m, 2H), 7.16 – 6.81 (m, 3H), 6.65 (d, *J* = 6.9 Hz, 1H), 6.11 (s, 2H). ^13^C NMR (101 MHz, DMSO-*d*_6_) δ 153.1, 148.8, 142.9, 141.5, 137.2, 136.5, 126.3, 122.6, 122.5, 120.0, 119.5, 119.1, 118.4, 107.9, 101.84, 101.80. HRMS *m*/*z* [M+H]^+^ calcd for C_16_H_12_N_2_O_2_Br: 343.0082, found 343.0072, LC *t*_R_ = 3.43 min, > 98 % Purity.

**6-Bromo-*N*-(2,5-difluorophenyl)quinolin-4-amine (10)** was obtained as a colourless solid (125 mg, 0.373 mmol, 60 %). MP 187-189 °C; ^1^H NMR (400 MHz, DMSO-*d*_6_) δ 11.11 (s, 1H), 9.21 (d, *J* = 2.0 Hz, 1H), 8.59 (d, *J* = 6.9 Hz, 1H), 8.20 (dd, *J* = 9.1, 1.9 Hz, 1H), 8.12 (d, *J* = 9.0 Hz, 1H), 7.78 – 7.46 (m, 2H), 7.38 – 7.25 (m, 1H), 6.56 (dd, *J* = 6.9, 2.3 Hz, 1H). ^13^C NMR (101 MHz, DMSO-*d*_6_) δ 161.5 (dd, *J* = 247.9, 11.7 Hz), 157.1 (dd, *J* = 251.6, 13.2 Hz), 154.5, 143.4, 137.3, 136.7, 130.2 (dd, *J* = 10.2, 2.0 Hz), 126.2, 122.7, 121.0 (dd, *J* = 12.6, 3.9 Hz), 120.2, 118.4, 113.0 (dd, *J* = 22.6, 3.7 Hz), 105.8 (dd, *J* = 27.1, 24.0 Hz), 100.9 (d, *J* = 1.8 Hz). HRMS *m*/*z* [M+H]^+^ calcd for C_15_H_10_N_2_F_2_Br: 334.9995, found 334.9985, LC *t*_R_ = 3.65 min, > 98% Purity.

**6-Bromo-*N*-(2,4-difluorophenyl)quinolin-4-amine (11)** was obtained as a colourless solid (94 mg, 0.281 mmol, 45 %). MP 301-303 °C; ^1^H NMR (400 MHz, DMSO-*d*_6_) δ 11.18 (s, 1H), 9.20 (d, *J* = 2.0 Hz, 1H), 8.62 (d, *J* = 6.8 Hz, 1H), 8.20 (dd, *J* = 9.0, 2.0 Hz, 1H), 8.12 (d, *J* = 9.0 Hz, 1H), 7.65 – 7.48 (m, 2H), 7.40 (ddt, *J* = 9.2, 8.0, 3.5 Hz, 1H), 6.68 (dd, *J* = 6.8, 2.7 Hz, 1H). ^13^C NMR (101 MHz, DMSO-*d*_6_) δ 158.2 (dd, *J* = 242.3, 2.3 Hz), 153.2 (dd, *J* = 245.2, 2.9 Hz), 153.9, 143.6, 137.4, 136.7, 126.2, 125.6 (dd, *J* = 14.7, 11.0 Hz), 122.8, 120.3, 118.5, 118.4 (dd, *J* = 22.5, 9.7 Hz), 116.3 (dd, *J* = 23.9, 8.2 Hz), 115.5, 115.2, 101.5 (d, *J* = 2.3 Hz). HRMS *m*/*z* [M+H]^+^ calcd for C_15_H_10_N_2_F_2_Br: 334.9995, found 334.9985, LC *t*_R_ = 3.63 min, > 98% Purity.

**6-Bromo-*N*-(perfluorophenyl)quinolin-4-amine (12)** was obtained as a brown solid(51.4 mg, 0.136 mmol, 21 %) MP 109-112 °C; ^1^H NMR (400 MHz, DMSO-*d*_6_) δ. ^13^C NMR (101 MHz, DMSO-*d*_6_) δ. HRMS *m*/*z* [M+H]^+^ calcd for C_16_H_7_N_2_F_8_: 379.0481, found 379.0473, LC *t*_R_ = 4.38 min, > 98% Purity.

**6-Bromo-*N*-(5-fluorobenzo[*d*][1,3]dioxol-4-yl)quinolin-4-amine (14)** was obtained as a beige solid (174 mg, 0.483 mmol, 78 %). MP 224-227 °C; ^1^H NMR (400 MHz, DMSO-*d*_6_) δ 11.08 (s, 1H), 9.26 (d, *J* = 1.7 Hz, 1H), 8.63 (d, *J* = 6.8 Hz, 1H), 8.28 – 8.06 (m, 2H), 7.04 (dd, *J* = 8.7, 4.1 Hz, 1H), 6.94 (dd, *J* = 10.8, 8.6 Hz, 1H), 6.67 (dd, *J* = 6.8, 1.2 Hz, 1H), 6.17 (s, 2H). ^13^C NMR (100 MHz, DMSO-*d*_6_) δ 153.8, 152.4 (d, *J* = 240.7 Hz), 144.8 (d, *J* = 1.8 Hz), 143.9 (d, *J* = 6.3 Hz), 143.4, 137.3, 136.8, 126.1, 122.7, 120.4, 118.4, 108.7 (d, *J* = 20.5 Hz), 107.9 (d, *J* = 22.4 Hz), 107.5 (d, *J* = 9.2 Hz), 103.2, 101.7. HRMS *m*/*z* [M+H]^+^ calcd for C_16_H_10_N_2_O_2_FBr: 359.9910, found 360.9987, LC *t*_R_ = 3.53 min, > 98 % Purity.

**6-Bromo-*N*-(2,3-dimethoxyphenyl)quinolin-4-amine (15)** was obtained as a colourless solid (187 mg, 0.520 mmol, 84 %). MP 150-153 °C; ^1^H NMR (400 MHz, DMSO-*d*_6_) δ 11.03 – 10.85 (br s, 1H), 9.19 (d, *J* = 2.0 Hz, 1H), 8.50 (d, *J* = 6.9 Hz, 1H), 8.18 (dd, *J* = 9.0, 2.0 Hz, 1H), 8.08 (d, *J* = 9.0 Hz, 1H), 7.26 (dd, *J* = 8.4, 7.7 Hz, 1H), 7.20 (dd, *J* = 8.4, 1.7 Hz, 1H), 7.00 (dd, *J* = 7.7, 1.7 Hz, 1H), 6.39 (d, *J* = 6.9 Hz, 1H), 3.89 (s, 3H), 3.67 (s, 3H). ^13^C NMR (101 MHz, DMSO-*d*_6_) δ 155.1, 154.0, 144.8, 143.2, 137.7, 137.0, 130.5, 126.6, 125.2, 123.0, 120.3, 119.9, 118.6, 113.5, 101.6, 61.2, 56.5. HRMS *m*/*z* [M+H]^+^ calcd for C_17_H_16_N_2_O_2_Br: 359.0395, found 359.0284, LC *t*_R_ = 3.62 min, > 98% Purity.

**6-Bromo-*N*-(2,2-difluorobenzo[*d*][1,3]dioxol-4-yl)quinolin-4-amine (16)** was obtained as a colourless solid (190 mg, 0.501 mmol, 81 %). MP 272-274 °C; ^1^H NMR (400 MHz, DMSO-*d*_6_) δ 11.36 (s, 1H), 9.22 (d, *J* = 2.0 Hz, 1H), 8.68 (d, *J* = 6.8 Hz, 1H), 8.21 (dd, *J* = 9.0, 2.0 Hz, 1H), 8.13 (d, *J* = 9.0 Hz, 1H), 7.50 (dd, *J* = 7.5, 1.7 Hz, 1H), 7.44 – 7.21 (m, 2H), 6.82 (d, *J* = 6.8 Hz, 1H). ^13^C NMR (100 MHz, DMSO-*d*_6_) δ 153.2, 144.1, 143.6, 137.4, 137.1, 136.8, 131.0 (t, *J* = 254.8 Hz), 126.3, 125.5, 122.7, 122.2, 120.4 (d, *J* = 3.0 Hz, 2C), 118.7, 109.2, 101.9. HRMS *m*/*z* [M+H]^+^ calcd for C_16_H_10_N_2_O_2_F_2_Br: 378.9894, found 378.9880, LC *t*_R_ = 4.13 min, > 98 % Purity.

**6-Bromo-*N*-(2,3-dihydrobenzofuran-4-yl)quinolin-4-amine (17)** was obtained as a yellow solid (133 mg, 0.390 mmol, 64 %). MP 291-293 °C; ^1^H NMR (400 MHz, DMSO-*d*_6_) δ 11.09 (s, 1H), 9.19 (d, *J* = 2.0 Hz, 1H), 8.52 (d, *J* = 6.9 Hz, 1H), 8.17 (dd, *J* = 9.0, 2.0 Hz, 1H), 8.08 (d, *J* = 9.0 Hz, 1H), 7.30 (t, *J* = 8.0 Hz, 1H), 6.89 (ddd, *J* = 10.4, 8.0, 0.8 Hz, 2H), 6.56 (d, *J* = 6.9 Hz, 1H), 4.59 (t, *J* = 8.7 Hz, 2H), 3.10 (t, *J* = 8.7 Hz, 2H). ^13^C NMR (100 MHz, DMSO-*d*_6_) 161.4, 153.6, 143.1, 137.3, 136.6, 133.7, 129.5, 126.3, 124.2, 122.5, 119.9, 118.5, 117.8, 108.6, 100.9, 71.2, 27.9. HRMS *m*/*z* [M+H]^+^ calcd for C_17_H_14_N_2_OBr: 341.0289, found 341.0283, LC *t*_R_ = 3.59 min, > 98 % Purity.

**6-Bromo-*N*-(2,3-dihydrobenzofuran-7-yl)quinolin-4-amine (18)** was obtained as a yellow solid (173 mg, 0.507 mmol, 82 %). MP 301-304 °C; ^1^H NMR (400 MHz, DMSO-*d*_6_) δ 11.08 (s, 1H), 9.25 – 9.17 (m, 1H), 8.52 (d, *J* = 6.9 Hz, 1H), 8.22 – 8.04 (m, 2H), 7.32 (dq, *J* = 7.3, 1.1 Hz, 1H), 7.23 – 7.15 (m, 1H), 6.99 (dd, *J* = 7.9, 7.3 Hz, 1H), 6.44 (d, *J* = 6.9 Hz, 1H), 4.59 (t, *J* = 8.7 Hz, 2H), 3.30 (t, *J* = 8.7 Hz, 2H). ^13^C NMR (101 MHz, DMSO-*d*_6_) δ 153.9, 153.5, 142.5, 137.1, 136.4, 129.8, 126.2, 125.5, 124.6, 122.4, 121.3, 119.8, 119.1, 118.2, 101.5, 71.9, 29.3. HRMS *m*/*z* [M+H]^+^ calcd for C_17_H_14_N_2_OBr: 341.0289, found 341.0285 LC *t*_R_ = 3.70 min, > 98 % Purity.

***N*-(Benzofuran-4-yl)-6-bromoquinolin-4-amine (19)** was obtained as a yellow solid (162 mg, 0.476 mmol, 77 %). MP >300 °C; ^1^H NMR (400 MHz, DMSO-*d*_6_) δ 11.36 (s, 1H), 9.27 (d, *J* = 2.0 Hz, 1H), 8.50 (d, *J* = 6.9 Hz, 1H), 8.20 (dd, *J* = 9.0, 2.0 Hz, 1H), 8.16 – 7.90 (m, 2H), 7.73 (dt, *J* = 8.3, 0.9 Hz, 1H), 7.51 (t, *J* = 8.0 Hz, 1H), 7.38 (dd, *J* = 7.7, 0.8 Hz, 1H), 6.91 (dd, *J* = 2.2, 1.0 Hz, 1H), 6.52 (d, *J* = 6.9 Hz, 1H). ^13^C NMR (100 MHz, DMSO-*d*_6_) δ 155.4, 154.0, 146.7, 143.0, 137.4, 136.5, 130.0, 129.7, 126.5, 125.3, 123.7, 122.5, 120.2, 119.9, 118.7, 111.1, 105.2, 101.0. HRMS *m*/*z* [M+H]^+^ calcd for C_17_H_12_N_2_OBr: 339.0133, found 339.0129, LC *t*_R_ = 3.82 min, > 98 % Purity.

***N*-(Benzofuran-7-yl)-6-bromoquinolin-4-amine (20)** was obtained as a yellow solid (157 mg, 0.464 mmol, 75 %). MP 256-259 °C; ^1^H NMR (400 MHz, DMSO-*d*_6_) δ 11.37 (s, 1H), 9.27 (d, *J* = 2.0 Hz, 1H), 8.53 (d, *J* = 6.9 Hz, 1H), 8.21 (dd, *J* = 9.1, 2.0 Hz, 1H), 8.13 (d, *J* = 9.0 Hz, 1H), 8.07 (d, *J* = 2.2 Hz, 1H), 7.79 (dd, *J* = 6.0, 2.9 Hz, 1H), 7.55 – 7.26 (m, 2H), 7.13 (d, *J* = 2.1 Hz, 1H), 6.46 (d, *J* = 6.9 Hz, 1H). ^13^C NMR (100 MHz, DMSO-*d*_6_) δ 154.0, 148.3, 146.7, 143.1, 137.3, 136.7, 129.5, 126.2, 124.0, 122.6, 122.0, 121.3, 121.2, 120.4, 118.5, 107.5, 101.4. HRMS *m*/*z* [M+H]^+^ calcd for C_17_H_12_N_2_OBr: 339.0133, found 339.0129, LC *t*_R_ = 4.36 min, > 98 % Purity.

***N*-(6-Bromoquinolin-4-yl)benzo[*d*]isoxazol-7-amine (21)** was obtained as a colourless solid (141 mg, 0.414 mmol, 67 %). MP >300 °C; ^1^H NMR (400 MHz, DMSO-*d*_6_) δ 11.49 (s, 1H), 9.45 (s, 1H), 9.29 (d, *J* = 2.0 Hz, 1H), 8.45 (d, *J* = 6.9 Hz, 1H), 8.31 (dd, *J* = 6.8, 2.4 Hz, 1H), 8.21 (dd, *J* = 9.0, 2.0 Hz, 1H), 8.11 (d, *J* = 9.0 Hz, 1H), 7.75 – 7.62 (m, 2H), 6.41 (d, *J* = 6.9 Hz, 1H). ^13^C NMR (100 MHz, DMSO-*d*_6_) δ 157.6, 154.5, 148.3, 142.7, 137.3, 136.6, 135.8, 130.7, 126.4, 126.2, 123.9, 122.6, 122.5, 120.0, 118.4, 101.9. HRMS *m*/*z* [M+H]^+^ calcd for C_16_H_11_N_3_OBr: 340.0085, found 340.0074, LC *t*_R_ = 3.18 min, > 98 % Purity.

***N*-(6-Bromoquinolin-4-yl)benzo[d]isoxazol-4-amine (22)** was obtained as a dark green solid (122 mg, 0.359 mmol, 58 %). MP >300 °C; ^1^H NMR (400 MHz, DMSO-*d*_6_) δ 11.37 (s, 1H), 9.31 (s, 1H), 9.24 (d, *J* = 2.0 Hz, 1H), 8.58 (d, *J* = 6.8 Hz, 1H), 8.22 (dd, *J* = 9.0, 1.9 Hz, 1H), 8.12 (d, *J* = 9.0 Hz, 1H), 7.93 – 7.70 (m, 2H), 7.53 (dd, *J* = 4.6, 3.7 Hz, 1H), 6.85 (d, *J* = 6.8 Hz, 1H). ^13^C NMR (101 MHz, DMSO-*d*_6_) δ 162.7, 153.9, 145.7, 143.7, 137.6, 136.7, 131.8, 131.4, 126.5, 122.8, 120.2, 120.1, 119.2, 117.4, 108.6, 101.6. HRMS *m*/*z* [M+H]^+^ calcd for C_16_H_11_N_3_OBr: 340.0085, found 340.0084, LC *t*_R_ = 3.38 min, > 98 % Purity.

***N*-(6-Bromoquinolin-4-yl)benzo[*d*]oxazol-7-amine (23)** was obtained as a dark yellow solid (65 mg, 0.192 mmol, 31 %). MP 305-308 °C; ^1^H NMR (400 MHz, DMSO-*d*_6_) δ 10.80 (s, 1H), 10.16 (s, 1H), 9.10 (d, *J* = 2.0 Hz, 1H), 8.58 (d, *J* = 6.9 Hz, 1H), 8.18 (dd, *J* = 9.0, 2.0 Hz, 1H), 8.03 (d, *J* = 9.0 Hz, 1H), 7.49 (d, *J* = 7.9 Hz, 1H), 7.21 (t, *J* = 7.9 Hz, 1H), 6.50 (d, *J* = 6.9 Hz, 1H). ^13^C NMR (176 MHz, DMSO) δ 160.9, 154.9, 149.2, 143.9, 143.3, 138.2, 136.8, 128.5, 127.0, 126.9, 126.3, 123.5, 121.1, 120.0, 119.1, 101.5. HRMS *m*/*z* [M+H]^+^ calcd for C_16_H_11_N_3_OBr: 340.0085, found 340.0084, LC *t*_R_ = 3.67 min, > 98 % Purity.

***N*-(6-Bromoquinolin-4-yl)benzo[*d*]thiazol-7-amine (24)** was obtained as a dark yellow solid (84 mg, 0.235 mmol, 38 %). MP 240-243 °C; ^1^H NMR (400 MHz, DMSO-*d*_6_) δ 9.54 (s, 1H), 9.39 (d, *J* = 0.4 Hz, 1H), 8.73 (t, *J* = 1.4 Hz, 1H), 8.44 (d, *J* = 5.2 Hz, 1H), 8.05 (dd, *J* = 8.2, 1.0 Hz, 1H), 7.96 – 7.75 (m, 2H), 7.66 (t, *J* = 7.9 Hz, 1H), 7.55 – 7.46 (m, 1H), 6.29 (d, *J* = 5.2 Hz, 1H). ^13^C NMR (100 MHz, DMSO-*d*_6_) δ 156.1, 154.9, 151.1, 147.5, 145.9, 134.7, 132.4, 131.5, 129.2, 127.3, 124.7, 121.8, 120.9, 120.4, 117.9, 103.6. HRMS *m*/*z* [M+H]^+^ calcd for C_16_H_11_N_3_F: 355.9857, found 355.9846, LC *t*_R_ = 3.20 min, > 98 % Purity.

***N*-(6-Bromoquinolin-4-yl)benzo[*d*]thiazol-4-amine (25)** was obtained as a colourless solid (150 mg, 0.421 mmol, 68 %). MP 281-283 °C; ^1^H NMR (400 MHz, DMSO-*d*_6_) δ 11.46 (s, 1H), 9.45 (s, 1H), 9.27 (d, *J* = 2.0 Hz, 1H), 8.45 (d, *J* = 6.9 Hz, 1H), 8.37 – 8.25 (m, 1H), 8.21 (dd, *J* = 9.0, 2.0 Hz, 1H), 8.10 (d, *J* = 9.0 Hz, 1H), 7.75 – 7.59 (m, 2H), 6.42 (d, *J* = 6.9 Hz, 1H). ^13^C NMR (100 MHz, DMSO-*d*_6_) δ 157.6, 154.4, 148.3, 142.7, 137.4, 136.6, 135.8 130.7, 126.4, 126.2, 123.9, 122.7, 122.5, 120.0, 118.4, 101.9. ^13^C NMR (101 MHz, DMSO-*d*_6_) δ. HRMS *m*/*z* [M+H]^+^ calcd for C_16_H_11_N_3_SBr: 355.9857, found 355.9846, LC *t*_R_ = 3.30 min, > 98 % Purity.

***N*-(Benzo[*b*]thiophen-4-yl)-6-bromoquinolin-4-amine (26)** was obtained as a dark yellow solid (147 mg, 0.414 mmol, 67 %). MP 308-311 °C; ^1^H NMR (400 MHz, DMSO-*d*_6_) δ 11.40 (s, 1H), 9.28 (d, *J* = 2.0 Hz, 1H), 8.46 (d, *J* = 6.9 Hz, 1H), 8.20 (dd, *J* = 9.0, 2.0 Hz, 1H), 8.16 (dt, *J* = 8.1, 0.9 Hz, 1H), 8.11 (d, *J* = 9.0 Hz, 1H), 7.88 (dd, *J* = 5.5, 0.4 Hz, 1H), 7.60 – 7.52 (m, 1H), 7.49 (dd, *J* = 7.5, 1.0 Hz, 1H), 7.37 (dd, *J* = 5.6, 0.8 Hz, 1H), 6.37 (d, *J* = 6.9 Hz, 1H). ^13^C NMR (100 MHz, DMSO-*d*_6_) δ 154.6, 143.0, 141.1, 137.4, 136.6, 135.8, 131.4, 129.1, 126.5, 125.3, 122.8, 122.5, 122.4, 121.3, 119.8, 118.6, 100.9. HRMS *m*/*z* [M+H]^+^ calcd for C_17_H_12_N_2_SBr: 354.9905, found 354.9900, LC *t*_R_ = 4.64 min, > 98 % Purity.

***N*-(1*H*-Benzo[*d*]imidazol-7-yl)-6-bromoquinolin-4-amine (27)** was obtained as a dark yellow solid (25 mg, 0.074 mmol, 12 %). MP 270-273 °C; ^1^H NMR (400 MHz, DMSO-*d*_6_) δ. ^13^C NMR (101 MHz, DMSO-*d*_6_) δ. HRMS *m*/*z* [M+H]^+^ calcd for C_16_H_12_N_4_Br: 339.0245, found 339.0232, LC *t*_R_ = 2.46 min, > 98% Purity.

***N*-(6-Bromoquinolin-4-yl)benzo[*c*][1,2,5]oxadiazol-4-amine (28)** was obtained as a black solid (152 mg, 0.445 mmol, 72 %). MP >300 °C; ^1^H NMR (400 MHz, DMSO-*d*_6_) δ 11.65 (s, 1H), 9.29 (d, *J* = 2.0 Hz, 1H), 8.63 (d, *J* = 6.8 Hz, 1H), 8.23 (dd, *J* = 9.0, 1.9 Hz, 1H), 8.18 – 8.06 (m, 2H), 7.83 – 7.70 (m, 2H), 6.94 (d, *J* = 6.8 Hz, 1H). ^13^C NMR (101 MHz, DMSO-*d*_6_) δ 153.6, 150.3, 145.9, 143.3, 137.4, 136.8, 133.2, 128.0, 126.4, 125.7, 122.8, 120.5, 118.9, 115.3, 103.5. HRMS *m*/*z* [M+H]^+^ calcd for C_15_H_10_N_4_OBr: 341.0038, found 341.0027, LC *t*_R_ = 3.39 min, > 98 % Purity.

***N*-(6-Bromoquinolin-4-yl)benzo[*c*][1,2,5]thiadiazol-4-amine (29)** was obtained as a yellow solid (164 mg, 0.458 mmol, 74 %). MP >300 °C; ^1^H NMR (400 MHz, DMSO-*d*_6_) δ 11.54 (s, 1H), 9.28 (d, *J* = 2.0 Hz, 1H), 8.48 (d, *J* = 6.8 Hz, 1H), 8.19 (ddd, *J* = 8.5, 6.4, 1.6 Hz, 2H), 8.12 (d, *J* = 9.0 Hz, 1H), 7.95 – 7.78 (m, 2H), 6.61 (d, *J* = 6.8 Hz, 1H). ^13^C NMR (100 MHz, DMSO-*d*_6_) δ 155.6, 153.8, 149.9, 143.3, 137.9, 136.5, 130.3, 129.3, 126.2, 126.1, 123.2, 120.8, 120.1, 118.8, 103.0. HRMS *m*/*z* [M+H]^+^ calcd for C_15_H_10_N_4_SBr: 356.9810, found 356.9798, LC *t*_R_ = 3.34 min, > 98 % Purity.

**6-Bromo-*N*-(1-methyl-1H-benzo[*d*][1,2,3]triazol-4-yl)quinolin-4-amine (30)** was obtained as a colourless solid (118 mg, 0.334 mmol, 54 %). MP 202-205 °C; ^1^H NMR (400 MHz, DMSO-*d*_6_) δ 9.54 (s, 1H), 8.82 (dd, *J* = 1.9, 0.8 Hz, 1H), 8.47 (d, *J* = 5.2 Hz, 1H), 7.98 – 7.75 (m, 2H), 7.64 (dd, *J* = 8.3, 0.9 Hz, 1H), 7.57 (dd, *J* = 8.3, 7.3 Hz, 1H), 7.31 (dd, *J* = 7.3, 1.0 Hz, 1H), 6.73 (d, *J* = 5.3 Hz, 1H), 4.33 (s, 3H). ^13^C NMR (100 MHz, DMSO-*d*_6_) δ 151.0, 147.5, 146.9, 139.6, 135.0, 132.2, 131.7, 131.4, 128.0, 124.9, 121.4, 117.9, 116.2, 106.2, 104.9, 34.3. HRMS *m*/*z* [M+H]^+^ calcd for C_16_H_13_N_5_Br: 354.0354, found 354.0343, LC *t*_R_ = 3.04 min, > 98 % Purity.

**6-Bromo-*N*-(1-methyl-1*H*-indazol-4-yl)quinolin-4-amine (31)** was obtained as a dark gray solid (131 mg, 0.371 mmol, 60 %). MP 206-209 °C; ^1^H NMR (400 MHz, DMSO-*d*_6_) δ 11.38 (s, 1H), 9.29 (d, *J* = 2.0 Hz, 1H), 8.49 (d, *J* = 6.9 Hz, 1H), 8.19 (dd, *J* = 9.0, 2.0 Hz, 1H), 8.12 (d, *J* = 9.0 Hz, 1H), 8.00 (d, *J* = 0.9 Hz, 1H), 7.75 (dt, *J* = 8.5, 0.8 Hz, 1H), 7.56 (dd, *J* = 8.5, 7.3 Hz, 1H), 7.26 (dd, *J* = 7.3, 0.7 Hz, 1H), 6.59 (d, *J* = 6.9 Hz, 1H), 4.12 (s, 3H). ^13^C NMR (101 MHz, DMSO-*d*_6_) δ 154.1, 142.9, 141.0, 137.4, 136.5, 130.9, 129.3, 126.8, 126.5, 122.5, 119.9, 119.2, 118.8, 117.2, 109.6, 101.3, 35.7. HRMS *m*/*z* [M+H]^+^ calcd for C_17_H_14_N_4_Br: 353.0402, found 353.0340, LC *t*_R_ = 3.48 min, > 98 % Purity.

**6-Bromo-*N*-(1*H*-indazol-4-yl)quinolin-4-amine (32)** was obtained as a bright yellow solid (174 mg, 0.513 mmol, 83 %). MP >300 °C; ^1^H NMR (400 MHz, DMSO-*d*_6_) δ 13.51 (s, 1H), 11.34 (s, 1H), 9.27 (d, *J* = 2.0 Hz, 1H), 8.50 (d, *J* = 6.9 Hz, 1H), 8.20 (dd, *J* = 9.0, 2.0 Hz, 1H), 8.11 (d, *J* = 9.0 Hz, 1H), 8.02 (d, *J* = 1.1 Hz, 1H), 7.65 (dt, *J* = 8.4, 0.9 Hz, 1H), 7.52 (dd, *J* = 8.4, 7.2 Hz, 1H), 7.23 (dd, *J* = 7.3, 0.7 Hz, 1H), 6.60 (d, *J* = 6.9 Hz, 1H). ^13^C NMR (100 MHz, DMSO-*d*_6_) δ 154.1, 143.0, 141.4, 137.4, 136.5, 131.9, 129.2, 126.8, 126.4, 122.5, 119.9, 118.8, 118.6, 117.1, 110.1, 101.2. HRMS *m*/*z* [M+H]^+^ calcd for C_16_H_12_N_4_Br: 339.0245, found 339.0234, LC *t*_R_ = 3.08 min, > 98 % Purity.

***N*-(1*H*-Indazol-4-yl)-6-(trifluoromethyl)quinolin-4-amine (33)** was obtained as a dark yellow solid (36 mg, 0.110 mmol, 17 %). MP 106-109 °C; ^1^H NMR (700 MHz, DMSO-*d*_6_) δ 9.27 (d, *J* = 4.6 Hz, 1H), 8.89 (s, 1H), 8.82 (d, *J* = 2.0 Hz, 1H), 8.41 (d, *J* = 8.9 Hz, 1H), 8.15 (dd, *J* = 8.9, 2.1 Hz, 1H), 8.02 (d, *J* = 4.6 Hz, 1H), 6.99 (d, *J* = 8.3 Hz, 1H), 6.93 (dd, *J* = 8.3, 7.1 Hz, 1H), 6.40 (d, *J* = 7.0 Hz, 1H), 5.51 (s, 1H), 3.50 (s, 1H). ^13^C NMR (176 MHz, DMSO-*d*_6_) δ 154.0, 150.6, 144.4, 138.8, 131.7, 128.1 (q, *J* = 32.1 Hz), 126.7, 126.6, 126.1, 125.2, 123.6, 123.4, 121.7, 118.4, 107.4, 104.8, 40.3. HRMS *m*/*z* [M+H]^+^ calcd for C_17_H_12_N_4_F_3_: 329.1014, found 353.0340, LC *t*_R_ = 5.66 min, > 98 % Purity.

***N*-(1*H*-Indazol-7-yl)-6-(trifluoromethyl)quinolin-4-amine (34)** was obtained as a bright yellow solid (189 mg, 0.576 mmol, 89 %). MP 251-254 °C; ^1^H NMR (400 MHz, DMSO-*d*_6_) δ 13.43 (s, 1H), 11.89 – 11.60 (m, 1H), 9.55 – 9.41 (m, 1H), 8.56 (d, *J* = 7.0 Hz, 1H), 8.47 – 8.21 (m, 2H), 8.15 (d, *J* = 1.0 Hz, 1H), 7.93 (dd, *J* = 8.5, 0.7 Hz, 1H), 7.70 (dt, *J* = 1.9, 0.9 Hz, 1H), 7.21 (dd, *J* = 8.5, 1.8 Hz, 1H), 6.89 (d, *J* = 7.0 Hz, 1H). ^13^C NMR (100 MHz, DMSO-*d*_6_) δ 156.2, 144.4, 140.6, 135.2, 134.1 (1C, br s), 129.7 (1C, d, *J* = 3.1 Hz), 128.3, 127.2 (q, *J* = 32.9 Hz), 125.6, 123.3 (d, *J* = 4.2 Hz), 122.8, 122.5, 122.3 (d, *J* = 5.9 Hz), 118.6, 117.2, 107.4 (1C, br s), 101.6. HRMS *m*/*z* [M+H]+ calcd for C_17_H_12_N_4_F_3_: 329.1014, found 329.1006, LC *t*_R_ = 3.42 min, > 98 % Purity.

**6-Bromo-*N*-(1*H*-indazol-6-yl)quinolin-4-amine (35)** was obtained as a orange solid (180 mg, 0.532 mmol, 86 %). MP >300 °C; ^1^H NMR (400 MHz, DMSO-*d*_6_) δ 13.34 (s, 1H), 11.19 (s, 1H), 9.18 (d, *J* = 2.0 Hz, 1H), 8.51 (d, *J* = 6.9 Hz, 1H), 8.47 – 8.09 (m, 2H), 8.07 (d, *J* = 9.0 Hz, 1H), 7.95 (dd, *J* = 8.5, 0.7 Hz, 1H), 7.66 (dt, *J* = 1.9, 0.9 Hz, 1H), 7.20 (dd, *J* = 8.5, 1.8 Hz, 1H), 6.88 (d, *J* = 6.9 Hz, 1H). ^13^C NMR (101 MHz, DMSO-*d*_6_) δ 154.3, 143.1, 140.2, 137.4, 136.5, 135.0, 133.8, 126.2, 122.6, 122.1, 121.9, 119.8, 118.7, 118.2, 106.7, 100.6. HRMS *m*/*z* [M+H]^+^ calcd for C_16_H_12_N_4_Br: 339.0245, found 339.0243, LC *t*_R_ = 3.33 min, > 98 % Purity.

**6-Bromo-*N*-(5-fluoro-1*H*-indazol-6-yl)quinolin-4-amine (36)** was obtained as a beige solid (188 mg, 0.526 mmol, 85 %). MP >300 °C; ^1^H NMR (400 MHz, DMSO-*d*_6_) δ 13.51 (s, 1H), 11.24 (s, 1H), 9.23 (d, *J* = 2.0 Hz, 1H), 8.56 (d, *J* = 6.9 Hz, 1H), 8.49 – 8.13 (m, 2H), 8.11 (d, *J* = 9.0 Hz, 1H), 7.88 (d, *J* = 10.3 Hz, 1H), 7.80 (dd, *J* = 6.3, 1.1 Hz, 1H), 6.61 (dd, *J* = 6.9, 2.5 Hz, 1H). ^13^C NMR (100 MHz, DMSO-*d*_6_) δ 154.8, 151.9 (d, *J* = 240.3 Hz), 143.3, 137.3, 136.7, 136.5, 134.4 – 133.0 (m, 1C), 126.2, 123.9 (d, *J* = 17.7 Hz), 122.6, 121.6 (d, *J* = 9.5 Hz), 120.1, 118.4, 109.8, 106.5 (d, *J* = 22.1 Hz), 101.2. HRMS *m*/*z* [M+H]^+^ calcd for C_16_H_11_N_4_FBr: 357.0151, found 357.0150, LC *t*_R_ = 3.43 min, > 98% Purity.

**6-Bromo-*N*-(1*H*-indazol-5-yl)quinolin-4-amine (37)** was obtained as a mustard solid (172 mg, 0.507 mmol, 82 %). MP 260-263 °C; ^1^H NMR (400 MHz, Methanol-*d*_4_) δ 8.89 (d, *J* = 2.0 Hz, 1H), 8.35 (d, *J* = 7.0 Hz, 1H), 8.28 – 8.00 (m, 2H), 8.00 – 7.81 (m, 2H), 7.76 (dt, *J* = 8.8, 0.9 Hz, 1H), 7.46 (dd, *J* = 8.8, 1.9 Hz, 1H), 6.82 (d, *J* = 7.0 Hz, 1H). ^13^C NMR (101 MHz, Methanol-*d*_4_) δ 157.2, 143.8, 140.8, 138.8, 138.3, 135.4, 131.1, 126.9, 126.1, 124.9, 123.3, 122.0, 120.0, 119.3, 113.1, 101.6. HRMS *m*/*z* [M+H]^+^ calcd for C_16_H_12_N_4_Br: 339.0245, found 339.0243, LC *t*_R_ = 3.18 min, > 98 % Purity.

**6-[(6-Bromoquinolin-4-yl)amino]-2,3-dihydro-1*H*-isoindol-1-one (38)** was obtained as a yellow solid (160 mg, 0.452 mmol, 73 %). MP >300 °C; ^1^H NMR (400 MHz, DMSO-*d*_6_) δ 11.19 (s, 1H), 9.18 (d, *J* = 2.0 Hz, 1H), 8.77 (s, 8.0, 2.0 Hz, 1H), 6.85 (d, *J* = 6.9 Hz, 1H), 4.45 (s, 2H). ^1^H NMR (400 MHz, DMSO-*d*_6_) δ 169.0, 154.0, 143.3, 142.9, 137.5, 136.9, 136.5, 134.3, 128.4, 126.2, 125.4, 122.7, 119.9, 119.4, 118.8, 100.5, 45.0. HRMS *m*/*z* [M+H]^+^ calcd for C_17_H_13_N_3_OBr: 354.0242, found 354.0240, LC *t*_R_ = 3.46 min, > 98 % Purity.

**5-[(6-Bromoquinolin-4-yl)amino]-2,3-dihydro-1*H*-1,3-benzodiazol-2-one (39)** was obtained as a yellow solid (180 mg, 0.507 mmol, 82 %). MP >300 °C; ^1^H NMR (400 MHz, DMSO-*d*_6_) δ 10.88 (p, J = 2.4 Hz, 3H), 9.07 (d, *J* = 2.1 Hz, 1H), 8.46 (d, *J* = 7.0 Hz, 1H), 8.15 (dd, *J* = 9.0, 2.0 Hz, 1H), 8.00 (d, *J* = 9.0 Hz, 1H), 7.08 (d, *J* = 8.7 Hz, 1H), 7.04 – 6.96 (m, 2H), 6.73 (d, *J* = 7.0 Hz, 1H). ^13^C NMR (100 MHz, DMSO-*d*_6_) δ 155.4, 154.5, 143.0, 137.5, 136.4, 130. 7, 129.7, 129.1, 126.0, 122.7, 119.6, 118.4, 117.9, 109.2, 106.1, 100.3. HRMS *m*/*z* [M+H]^+^ calcd for C_16_H_12_N_4_OBr: 355.0194, found 355.0195, LC *t*_R_ = 3.43 min, > 98 % Purity.

**6-[(6-Bromoquinolin-4-yl)amino]-2,3-dihydro-1,3-benzoxazol-2-one (40)** was obtained as a yellow solid (165 mg, 0.464 mmol, 75 %). MP >300 °C; ^1^H NMR (400 MHz, Methanol-*d*_4_) δ 8.84 (d, *J* = 1.9 Hz, 1H), 8.40 (d, *J* = 6.9 Hz, 1H), 8.14 (dd, *J* = 9.0, 2.0 Hz, 1H), 7.87 (d, *J* = 9.0 Hz, 1H), 7.40 (dd, *J* = 1.7, 0.8 Hz, 1H), 7.35 – 7.18 (m, 2H), 6.89 (d, *J* = 6.9 Hz, 1H). ^13^C NMR (101 MHz, Methanol-*d*_4_) δ 156.9, 156.5, 146.1, 144.6, 138.1, 132.8, 131.6, 126.8, 123.9, 123.8, 122.9, 122.0, 120.2, 111.7, 109.0, 101.9. HRMS *m*/*z* [M+H]^+^ calcd for C_16_H_11_N_3_O_2_Br: 356.0035, found 356.0032, LC *t*_R_ = 3.66 min, > 98 % Purity.

**5-[(6-Bromoquinolin-4-yl)amino]-2,3-dihydro-1,3-benzoxazol-2-one (41)** was obtained as a yellow solid (163 mg, 0.458 mmol, 74 %). MP >300 °C; ^1^H NMR (400 MHz, DMSO-*d*_6_) δ 12.01 (s, 1H), 11.07 (s, 1H), 9.14 (d, *J* = 2.0 Hz, 1H), 8.51 (d, *J* = 6.9 Hz, 1H), 8.16 (dd, *J* = 9.0, 2.0 Hz, 1H), 8.05 (d, *J* = 9.0 Hz, 1H), 7.46 (d, *J* = 8.4 Hz, 1H), 7.30 – 6.99 (m, 2H), 6.79 (d, *J* = 6.9 Hz, 1H). ^13^C NMR (100 MHz, DMSO-*d*_6_) δ 154.5, 154.4, 143.1, 142.3, 137.4, 136.5, 132.8, 131.6, 126.2, 122.5, 119.8, 119.2, 118.5, 110.5, 107.5, 100.5. HRMS *m*/*z* [M+H]^+^ calcd for C_16_H_11_N_3_O_2_Br: 356.0035, found 356.0032, LC *t*_R_ = 3.71 min, > 98 % Purity.

**5-[(6-Bromoquinolin-4-yl)amino]-1-methyl-2,3-dihydro-1*H*-1,3-benzodiazol-2-one (42)** was obtained as a yellow solid (162 mg, 0.439 mmol, 71 %). MP >300 °C; ^1^H NMR (400 MHz, DMSO-*d*_6_) δ 11.16 (s, 1H), 11.10 (s, 1H), 9.17 (d, *J* = 2.0 Hz, 1H), 8.46 (d, *J* = 7.0 Hz, 1H), 8.13 (dd, *J* = 9.0, 1.9 Hz, 1H), 8.05 (d, *J* = 9.0 Hz, 1H), 7.23 (d, *J* = 8.2 Hz, 1H), 7.17 – 6.98 (m, 2H), 6.71 (d, *J* = 7.0 Hz, 1H), 3.33 (s, 3H). ^13^C NMR (100 MHz, DMSO-*d*_6_) δ 154.6, 154.5, 142.7, 137.3, 136.4, 130.3, 130.1, 129.1, 126.2, 122.4, 119.6, 118.4, 118.0, 108.3, 106.3, 100.2, 26.6. HRMS *m*/*z* [M+H]^+^ calcd for C_17_H_14_N_4_OBr: 369.0351, found 369.0350 LC *t*_R_ = 3.70 min, > 98 % Purity.

**5-((6-Bromoquinolin-4-yl)amino)-1,3-dihydrobenzo[*c*]thiophene 2,2-dioxide (43)** was obtained as an orange solid (205 mg, 0.526 mmol, 85 %). MP >300 °C; ^1^H NMR (400 MHz, DMSO-*d*_6_) δ 11.19 (s, 1H), 9.19 (d, *J* = 2.0 Hz, 1H), 8.55 (d, *J* = 6.9 Hz, 1H), 8.17 (dd, *J* = 9.0, 1.9 Hz, 1H), 8.09 (d, *J* = 9.0 Hz, 1H), 7.58 (d, *J* = 8.2 Hz, 1H), 7.54 – 7.36 (m, 2H), 6.88 (d, *J* = 6.9 Hz, 1H), 4.59 (d, *J* = 7.1 Hz, 4H). ^13^C NMR (100 MHz, DMSO-*d*_6_) δ 153.9, 143.2, 137.4, 136.9 136.6, 134.1, 131.5, 127.5, 126.3, 125.0, 122.5, 120.0, 118.7, 100.5, 56.1, 55.8. HRMS *m*/*z* [M+H]^+^ calcd for C_17_H_14_N_2_O_2_SBr: 388.9959, found 388.9955, LC *t*_R_ = 3.65min, > 98 % Purity.

## ACKNOWLEDGMENT

The SGC is a registered charity (number 1097737) that receives funds from AbbVie, Bayer Pharma AG, Boehringer Ingelheim, Canada Foundation for Innovation, Eshelman Institute for Innovation, Genome Canada, Innovative Medicines Initiative (EU/EFPIA) [ULTRA-DD grant no. 115766], Janssen, Merck KGaA Darmstadt Germany, MSD, Novartis Pharma AG, Ontario Ministry of Economic Development and Innovation, Pfizer, São Paulo Research Foundation-FAPESP, Takeda, and Wellcome [106169/ZZ14/Z]. We thank Biocenter Finland/DDCB for financial support and the CSC-IT Center for Science Ltd. (Finland) for allocation of computational resources. We acknowledge Dr. Brandie Ehrmann for LC-MS/HRMS support provided by the Mass Spectrometry Core Laboratory at the University of North Carolina at Chapel Hill. We are thankful for useful discussions with Antti Poso and Minna Rahnasto (University of Eastern Finland).

## ASSOCIATED CONTENT

**Supporting Information**. Experimental procedures for the synthesis of intermediates together with biological data and associated errors, NMR conformation and molecular formula strings (PDF) Molecular formula strings (CSV)

## Author Contributions

The manuscript was written through contributions of all authors. All authors approved of the final version of the manuscript.

